# Continuous nucleolar ribosomal RNA synthesis in differentiating lens fiber cells until abrupt nuclear degradation required for ocular lens transparency

**DOI:** 10.1101/2024.10.21.619434

**Authors:** Danielle Rayêe, U. Thomas Meier, Carolina Eliscovich, Aleš Cvekl

## Abstract

Cellular differentiation requires highly coordinate action of all three transcriptional systems to produce rRNAs, mRNAs, and various “short” and “long” non-coding RNAs by RNA Polymerase I, II, and III systems, respectively. The RNA Polymerase I catalyzes transcription of about 400 copies of rDNA genes generating 18S, 5.8S, and 28S rRNA molecules from the individual primary transcript. Lens fiber cell differentiation is a unique process to study transcriptional mechanisms of individual crystallin genes as their very high transcriptional outputs are directly comparable only to globin genes in erythrocytes. Importantly, both terminally differentiated lens fiber cells and mammalian erythrocytes degrade their nuclei though by different mechanisms. In lens, generation of organelle-free zone (OFZ) includes degradation of mitochondria, endoplasmic reticulum, Golgi apparatus, and nuclei; nevertheless, very little is known about their nucleoli and rRNA transcription. Here, using RNA fluorescence *in situ* hybridization (FISH) we evaluated nascent rRNA transcription during the entire process of lens fiber cell differentiation. The lens fiber cell nuclei undergo morphological changes prior their denucleation, including chromatin condensation; remarkably, the nascent rRNA transcription persists in all nuclei next to the OFZ. The changes in both nuclei and nucleoli shape and microarchitecture were evaluated by immunofluorescence to detect fibrillarin, nucleolin, UBF, and other nuclear proteins. These studies demonstrate for the first time that highly condensed lens fiber cell nuclei have the capacity to support rRNA transcription. Thus, “late” production of rRNA molecules and consequently the ribosomes contribute to the terminal translational mechanisms to produce maximal quantities of the crystallin proteins.

## INTRODUCTION

The eukaryotic cell nuclei operate three distinct transcriptional systems known as RNA polymerase I, II, and III (Roeder, 2019). The RNA Polymerase I system employs about 400 copies of rDNA genes in mammalian genomes to generate individual mature 18S, 5.8S, and 28S rRNA molecules from a single primary transcript (Feng and Manley, 2022) and is located within a single or a few nucleoli of the individual nucleus (Dundr and Misteli, 2001). The nucleoli (Olson et al., 2000; Boisvert et al., 2007; Correll et al., 2019) are membrane-less nuclear sub-compartments (Lafontaine et al., 2021; Yoneda et al., 2021) composed of the granular component (GC), the dense fibrillar component (DFC), and the fibrillar center (FC), and transcription occurs between the FC and DFC compartments (McStay, 2016; Hori et al., 2023). Following rRNA processing, the resulting mature rRNA molecules are used as individual structural and catalytical components of the large and small ribosomal subunits assembled in the nucleolus into the individual ribosomes (Granneman and Baserga, 2005). The rDNA genomic sequences (Zentner et al., 2014), located on short arms of acrocentric chromosomes (12, 15, 16, 17, 18, and 19 in mouse) represent the most evolutionarily conserved domains of the genome (Cerqueira and Lemos, 2019; Correll et al., 2019). In contrast, the RNA Polymerase II system generates mRNAs from over 23,000 protein coding genes as well as a growing list of lncRNAs (Cech and Steitz, 2016). Finally, the RNA Polymerase III system generates 5S rRNA, tRNAs, and various snRNAs, and plays a role in homologous repair of DNA double-stranded breaks (White, 2011). In an average mammalian cell, at the quantitative level, rRNAs and mRNAs represent 80-90% and ∼4% of the total RNA, respectively, with other operational RNAs (tRNAs, lncRNAs, snRNA, snoRNAs, eRNAs, and miRNAs) making up the rest (Wu et al., 2014; Feng and Manley, 2022). The 3-dimensional (3D) chromatin organization within the individual nuclei and its role in transcription of individual genes is cell type-specific and represents one of the most intricate biological puzzles to understand coordinated action of complex machineries regulating transcription, DNA replication, and DNA repair (Vermunt et al., 2019; Misteli, 2020).

Cell growth and proliferation are proportional to the rate of protein synthesis driven by ribosome biogenesis (Grummt and Ladurner, 2008; Ni and Buszczak, 2023). Individual cells invest major resources into the ribosome synthesis and these processes change in postmitotic terminally differentiated cells. However, there is a unique situation shared by mammalian erythrocytes and ocular lens fiber cells as they degrade their nuclei (Zhao et al., 2016; Limi et al., 2018). In lens, the nuclei located in the central subregion of the lens fiber cell compartment gradually change their shape and size prior their abrupt physical disintegration (Bassnett, 2009; Rowan et al., 2017; Brennan et al., 2019). This unique denucleation process is essential for lens transparency together with a robust accumulation of individual α- and β/γ-crystallin proteins, encoded by 16 genes of the mouse genome, in the lens fiber cell cytoplasm reaching concentration of 450 mg/ml (Cvekl and Eliscovich, 2021). Thus, the lens is an excellent system to examine molecular mechanism required for both tissue-specific and maximal transcriptional and translational outputs (Cvekl and Eliscovich, 2021).

We have shown earlier that expression of crystallin genes is only quantitatively comparable to the expression of globin genes in red blood cells while other highly expressed genes, such as insulins (Ins1 and Ins2) and α2-HS glycoprotein (Ahsg) and transferrin (Trf) are expressed in much lower levels in the pancreas and liver, respectively (Sun et al., 2015). Unexpectedly, our analysis of nascent mRNA expression of βA1- and γA-crystallin mRNAs revealed that their expression peaks in lens fiber cell nuclei just prior their destruction while transcription of the αA-crystallin genes generating the most abundant lens proteins peaked much earlier (Limi et al., 2018). Strikingly, nearly 60% of nuclei just prior their degradation showed expression of at least one γA-crystallin (*Cryga*) allele with biallelic expression found in more than 25% of nuclei. Visualization of transcriptionally active RNA Polymerase II enzymes found a few large condensates in these nuclei concomitant with active transcription of multiple crystallin genes (Limi et al., 2018). However, nothing is known about nascent transcription catalyzed by RNA Polymerase I within the lens fiber cell nucleoli.

To produce ribosomes, the RNA Polymerase I system (Grummt, 2003) requires the major portion of nutritionally-demanding resources (Grummt and Ladurner, 2008; Cerqueira and Lemos, 2019; Shore and Albert, 2022). However, the operations of lens fiber cell transcriptional-translational factories are limited by the denucleation process and reduced direct blood support. Degradation of mitochondria, endoplasmic reticulum (ER), and Golgi apparatus also proceeds in terminally differentiating lens fiber cells (Brennan et al., 2018). Another challenge for delivery of nutrients to the lens is degradation of the transient vascular systems located both at the anterior and posterior portions of the lens (McKeller et al., 2002; Chen et al., 2008; Beebe, 2008). Thus, it is possible that rRNA production may be attenuated earlier, reduced, or kept at high levels till the nuclear degradation depending how lens fiber cells evolved to manage their precious resources for generation of ribosomes and subsequently crystallin proteins (Cvekl and Eliscovich, 2021).

Earlier studies of the lens fiber cell nuclei revealed changes of nuclear shape from ovoid nuclei in lens epithelial cells and in early elongating lens fibers into more rounded nuclei (Bassnett, 2009). Like in mammalian erythrocytes (Zhao et al., 2016), these nuclei show chromatin condensation, transfer of histone and non-histone proteins into the cytoplasm, and reduction of their sizes; however, lens nuclei undergo an abrupt fragmentation within the cytoplasm (Limi et al., 2018). In contrast, the erythrocyte nuclei are extruded from the cells (Rogerson et al., 2018). Previous studies of lens fiber cell nucleoli and rRNA transcription were limited to general studies of nuclear morphology (Dahm et al., 1998; Gribbon et al., 2002) and detection of the 47S pre-rRNAs in proliferating cells of the adult mouse lens epithelium (Qian et al., 2006).

To generate new insights into the biology of the nucleolus during lens fiber cell differentiation, we analyzed various proteins located both in the nucleolus and nucleus together with RNA FISH to visualize nascent rRNA transcription and mature rRNAs during mouse lens embryogenesis. The results clearly show that nuclei next to the OFZ are capable of nascent rDNA transcription catalyzed by RNA Polymerase I, and, thus, retain functional nucleoli.

## RESULTS

### Visualization of nuclear morphology, fibrillarin and nucleolin through the progressive stages of mouse lens fiber cell differentiation

To follow the lens fiber cell differentiation, the lens fiber cell compartment of the newborn lens can be divided into four symmetric sub-regions a, b, c and d (Figs. 1A, B) to aid the initial spatial visualization of gradual changes in the nuclear morphology within the concentric zones of lens fiber cells at various stages of their terminal differentiation (Limi et al., 2018). Regions a, b, c and d represent postmitotic early differentiated fiber cells, intermediate fiber cells, advanced fiber cells, and pre-denucleated cells, respectively. At late embryonic stage E18.5, the OFZ is not yet fully established, and area d comprises secondary fiber cells at late differentiation phase (Fig. 1A). In the newborn lens, the OFZ is in the center of the tissue (Fig. 1B - region d). Nuclear morphology changes are visible throughout these individual stages of fiber cells differentiation. In area a, there are abundant oval shaped nuclei whereas areas b and c display more rounded nuclei undergoing chromatin condensation (Fig. 1C). In outer subregions of area d, small scattered nuclear traces are found (Fig. 1C).

**FIGURE 1.**
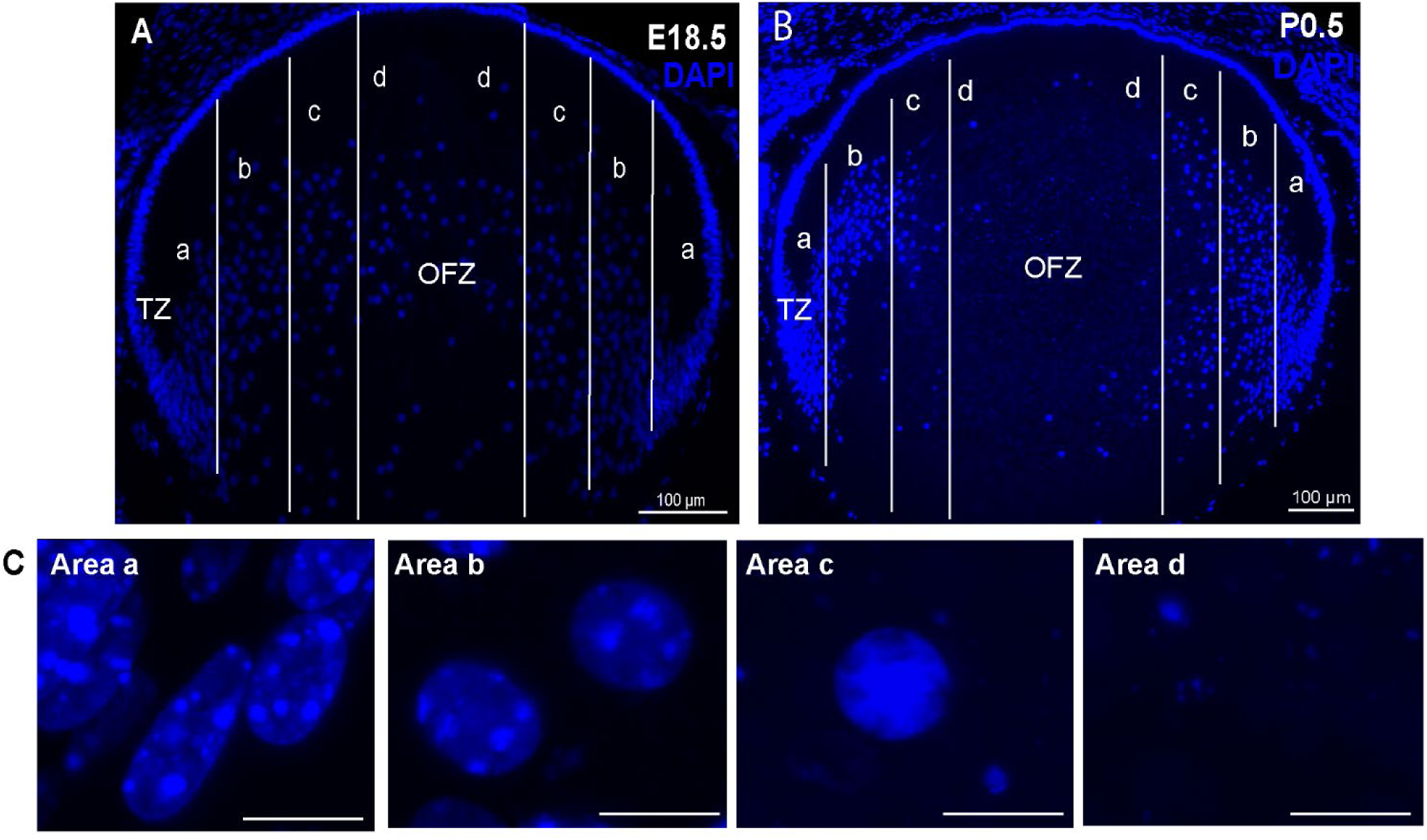
Mouse lens morphology and nuclear changes throughout denucleation. (A) The E18.5 lens is symmetrically divided into areas a, b, c and d from the transition zone (TZ, cells exiting cell cycle and undergoing the earliest stages of terminal differentiation) to the imminent organelle free zone (OFZ), yet not completely formed. (B) The P0.5 newborn lens display the OFZ within the area d, with no remaining nuclei in that area. (C) Inserts from areas A-D displaying nuclear morphological changes prior the denucleation. Area a is marked by elongated and abundant nuclei, area b shows intermediate fiber cells prior to the nuclear condensation in area c, and complete nuclear disintegration is found in the area d. Inserts scale bars = 10 µm. DAPI-stained nuclei (blue).

To explore how primary and secondary fiber cells change their nuclear and nucleolar morphology throughout their differentiation process, we first analyzed cells in areas a, b, c and d using immunofluorescence to visualize fibrillarin (rRNA 2’-O-methyltransferase, 34.3 kDa) as the canonical marker of the nucleolus, and lamin B1 of the inner nuclear membrane, at embryonic stages E14.5 and E16.5. At E14.5, the majority of lens fiber cell compartment is formed from the “primary” lens fiber cells that are formed from the posterior cells of the lens vesicle (E11-E11.5) via cell cycle exit-coupled terminal differentiation (Cvekl and Zhang, 2017). At E14.5, both the areas a and b are marked by two or three nucleoli per nucleus as observed by the fibrillarin stainings (Fig. 2A). The nuclear morphology is overall elongated throughout all areas a-d (Fig. 2B). At E16.5, secondary lens fiber cells also show elongated nuclei in areas a-b. In contrast, nuclei in areas c-d formed by primary lens fibers exhibit more rounded shape and marked nuclear condensation (Figs. 2C, D). These trends are further highlighted at E18.5 embryonic lenses which display stretched-shape nuclei found in areas a and b as well as abundant nucleoli staining (Figs. 3A, B). At E18.5, fiber cells in area d clearly show signs of nuclear condensation and nucleoli co-localizing with DAPI-negative nuclear regions (Figs. 3A, B). Finally, the newborn (P0.5) lenses have OFZ formed in area d (Fig. 3D). Fibrillarin stainings show that the cells in areas a and b display large and multiple nucleoli per their single nucleus. In contrast, some nuclei in the area c are highly condensed within a rounded shape; nevertheless, normal nucleoli stainings are present (Fig. 3C, D). With the advanced lens fiber cell differentiation, nuclear shape drastically shifts from elongated to round-shaped from area a to area b (Fig. 3D). Remarkably, we observed that the nuclear lamina remains intact in the denucleating cells within the area c (Fig. 3D).

**FIGURE 2.**
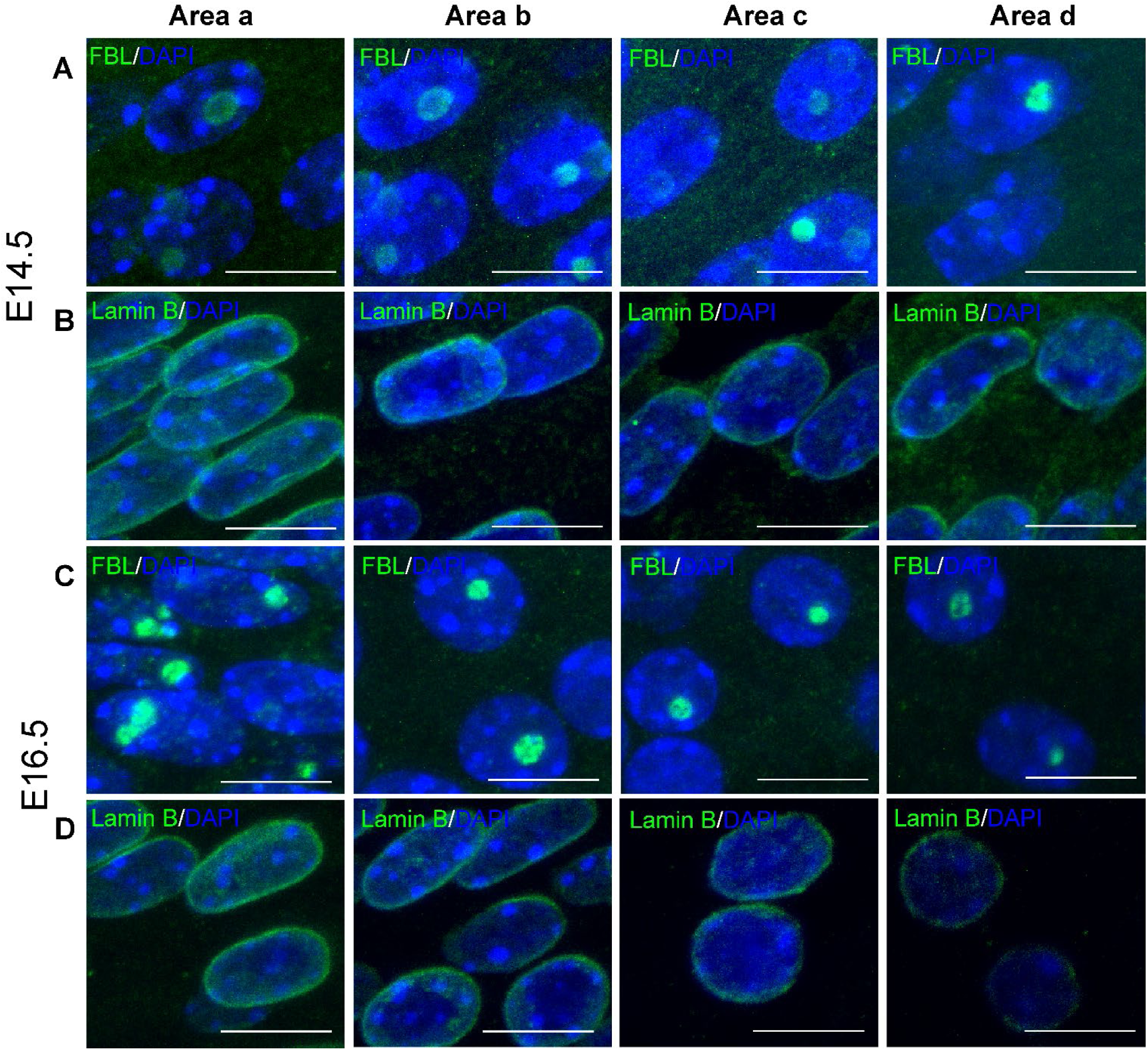
Early stages of differentiation of mouse lens fiber cells (E14.5 and E16.5). Immunolabeling of fibrillarin and lamin B (both in green) was performed to show primary (E14.5) and both primary and secondary (E16.5) lens fiber cells nuclei and nucleoli morphology in lens tissue regions (panels A-D). The segmented areas analyzed are shown in Fig. 1A. (A-D) Fiber cells in area a display consistently elongated nuclei throughout embryonic stages E14.5 to E16.5. (B) E14.5 lens is comprised mostly of primary lens fiber cells that display abundant nucleolar staining, pattern also seen in secondary fiber cells in E16.5. E14.5 nuclear morphology is mostly elongated whereas in E16.5 rounded-shape nuclei are seen more frequently in areas c and d. (C-D) E16.5 advanced primary lens fiber cells nuclei display nuclear condensation observed in DAPI and lamin B stainings. Scale bars = 10 µm. DAPI-stained nuclei (blue).

**FIGURE 3.**
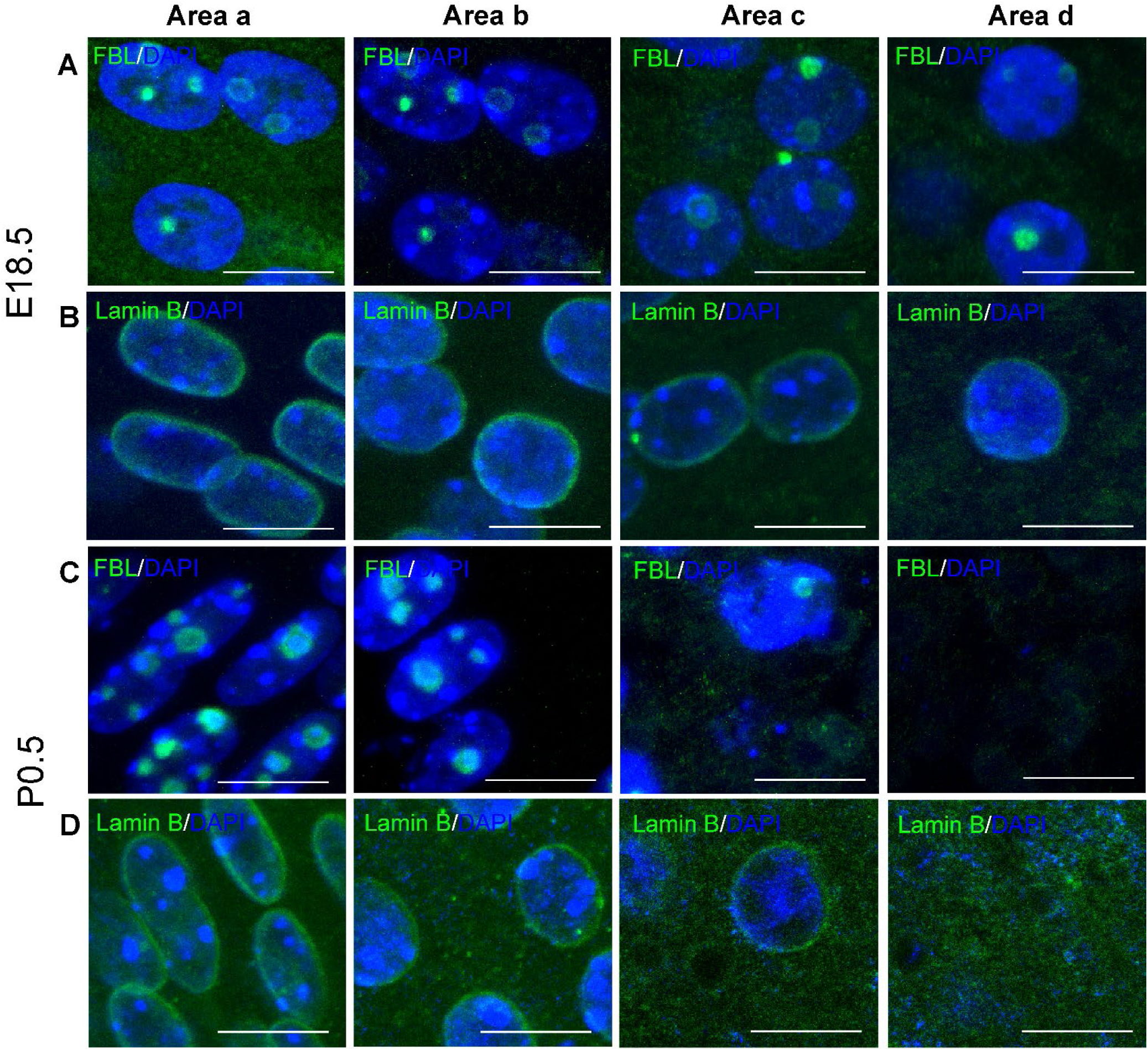
The final stages of lens fiber cells differentiation (E18.5 and P0.5) are marked by nuclear disassembly. (A-D) E18.5 and P0.5 lenses display elongated nuclei in area a, and several nucleoli co-localizing with fibrillarin. (A-B) In the area b, there is a mixed population of cells transitioning into the center of lens tissue. (A-D) In the area c fiber cells start displaying first signs of nuclear condensation at E18.5, however morphological changes are more obvious at newborn lens (bottom). (C) Nucleolar structure (green) remains intact in fiber cells at advanced stage of differentiation (white arrowheads in area c). Nuclear envelope (green) is observed in late degenerated nuclei in area c. (A,D) Sparse nuclei are found in the area d of E18.5 lens tissue whereas at P0.5 the OFZ is already formed. Scale bars = 10 µm. DAPI-stained nuclei (blue).

Nucleolin (77 kDa) is a major nucleolar protein, comprised of four RNA-binding domains and C-terminal RGG-rich “tail”, also located in the nucleoplasm known to play a role in rDNA transcription and ribosome assembly and their maturation (Scott and Oeffinger, 2016). Importantly, nucleolin co-localizes with the nucleoli during active transcription (Ma et al., 2007). We observed nucleolin signal co-localizing with the nucleolus throughout areas a-d in the mouse E14.5, E16.5 and E18.5 embryonic fiber cells (Fig. 4A, C). Nuclei displaying more than one nucleolus are frequently seen in all these four areas. Nevertheless, nucleolar numbers per each nucleus remain consistent throughout the embryonic and final stages of lens fiber cell differentiation (Supplementary Fig. S1A, B).

**FIGURE 4.**
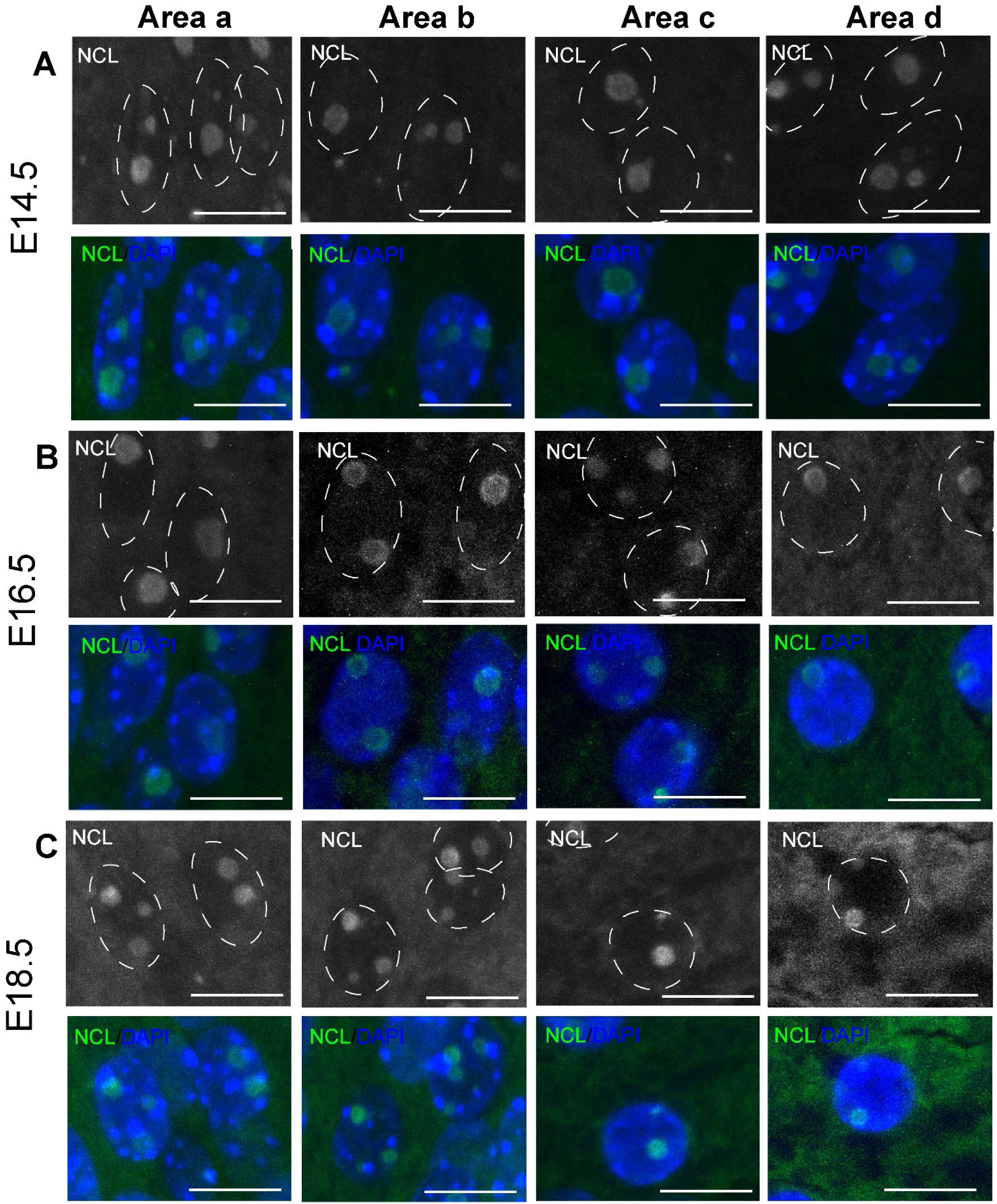
Nucleoli staining in embryonic lens fiber cells. The data show E14.5 (A), E16.5 (B) and E18.5 (C) embryonic stages in areas a, b, c and d. Nucleolin (NCL) and nuclei dotted (top); NCL (green) merged with DAPI-stained nuclei (bottom). Scale bars = 10 µm.

Previous studies of mouse and chick lenses demonstrated active DNA repair in maturing nuclei within the prospective and adjacent to the OFZ (Wang et al., 2010; Gheyas and Menko, 2023). To investigate this further, we immunolabelled newborn (P0.5) lens tissue for the histone variant H2AX phosphorylated at serine 139 residue (γH2AX) (Economopoulou et al., 2019). Nuclei displayed progressive increase in γH2AX staining in areas b and c (Fig. 5A). We next quantified the percentage of γH2AX cells throughout areas a-c. Areas b and c show a significant increase in the γH2AX positive nuclei per total nuclei in three biological replicates (Figure 5B). These data suggest that fiber cells approaching the OFZ start displaying signs of DNA damage and active repair in the area b. However, all cells in the area c exhibit increased nuclear damage/repair prior to the denucleation process. Taken together, there are gradual changes in the nuclear morphology including the number of nucleoli during lens fiber cell terminal differentiation that may negatively impact transcriptional processes to generate rRNAs.

**FIGURE 5.**
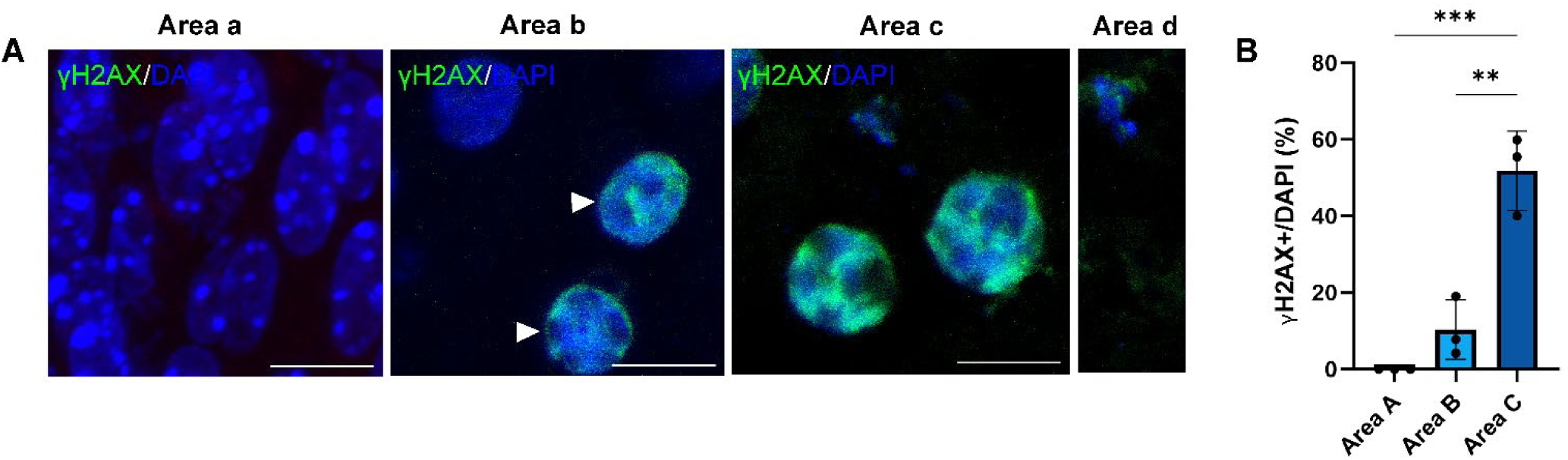
Analysis of DNA damage/repair prior the lens fiber cell denucleation process (P0.5). (A) The γH2AX positive nuclei are more frequently observed in areas b (white arrowheads) and c. DAPI-stained nuclei (blue). (B) Quantification of γH2AX positive nuclei per total number of nuclei throughout areas a-c. Statistical significance is shown by p-values; n=3 biological replicates. Scale bars = 10 µm.

### Nascent and mature rRNAs are generated in nucleoli adjacent to the OFZ

To establish rRNA FISH in the mouse lens, we employed two sets of probes targeting the rRNA internal-transcribed spacers 1 and 2 (ITS1/ITS2 probes) and the 18S and 28S regions (18S/28S probes) of the pre-rRNAs, as described elsewhere (Chebrout et al., 2022) (Fig. 6A). The ITS1/ITS2 probes localize the nascent nucleolar rRNAs, whereas the 18S/28S probes visualize mature pre-RNAs that accumulate in the nucleus and are already exported to the cytoplasm within the individual ribosomes.

**FIGURE 6.**
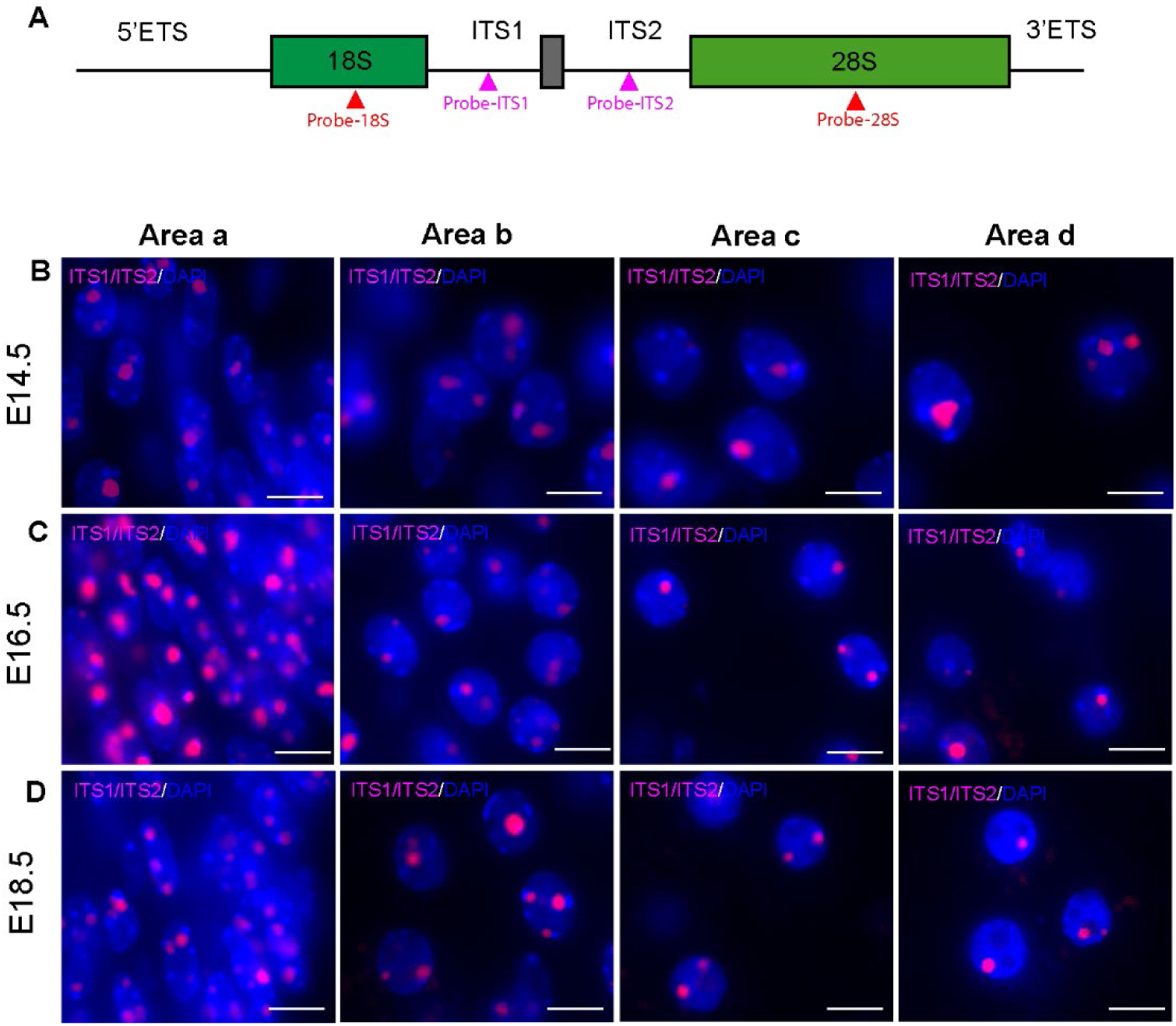
Visualization of nascent rRNA transcription throughout lens embryonic development (E14.5-E18.5). (A) A representative scheme of probes location in the internal-transcribed spacers 1 and 2 (ITS1/ITS2, magenta) and in the 18S and 28S (red) regions of the 47S pre-rRNAs. rRNA FISH *in situ* hybridization using ITS1/ITS2 (magenta) probes was performed in embryonic lens tissue sections. (A) E14.5 fiber cells display nucleolar rRNA signal throughout areas a-d. (B) E16.5 fiber cells in area a display various nucleoli rRNA signal per nucleus. (C) E18.5 fiber cells display rRNA nucleolar signal in all areas, including area d, where condensed nuclei are observed. 5’ETS: 5’external transcribed spacer; 3’ETS: 3’external transcribed spacer. Scale bars = 5 µm. Nuclei stained with DAPI (blue).

We first analyzed patterns of rRNA signals in embryonic lenses throughout development from E14.5 and E18.5 by *in situ* hybridization with ITS1/ITS2 probes (Figure 6A-C). We observed that fiber cells in area a show frequently more than one nucleolus per nucleus in all embryonic stages. Nucleolar rRNA follows similar patterns seen in nucleolar staining with NCL and fibrillarin (Figs. 3 and 4). At E18.5, area d, nuclei display the first signs of condensation (Fig. 6D).

Pre-rRNAs co-localizing with nucleoli in areas a and b in newborn lenses were observed (Fig. 7A, top; Fig. 7B, top). Importantly, the nuclei located in areas c and d also display nascent rRNAs in their nucleoli (Figure 7A, B top and bottom). It is possible to detect the ITS1/ITS2 signals even in highly condensed nuclei located in area c. In area d representing OFZ, no intranuclear signal was detected as expected. Mature rRNAs detected via the 18S/28S probes are abundant in the nuclei in areas a and b of the newborn mouse lens (Fig. 8A, bottom). Nuclei displaying more than one nucleolus were often observed in areas a and b. High cytoplasmic rRNA FISH signal was also detected using the 18S/28S probe set (Figs. 7A, B bottom). This data demonstrate that the mature rRNAs are present in the cytoplasm of lens fiber cells after their denucleation.

**FIGURE 7.**
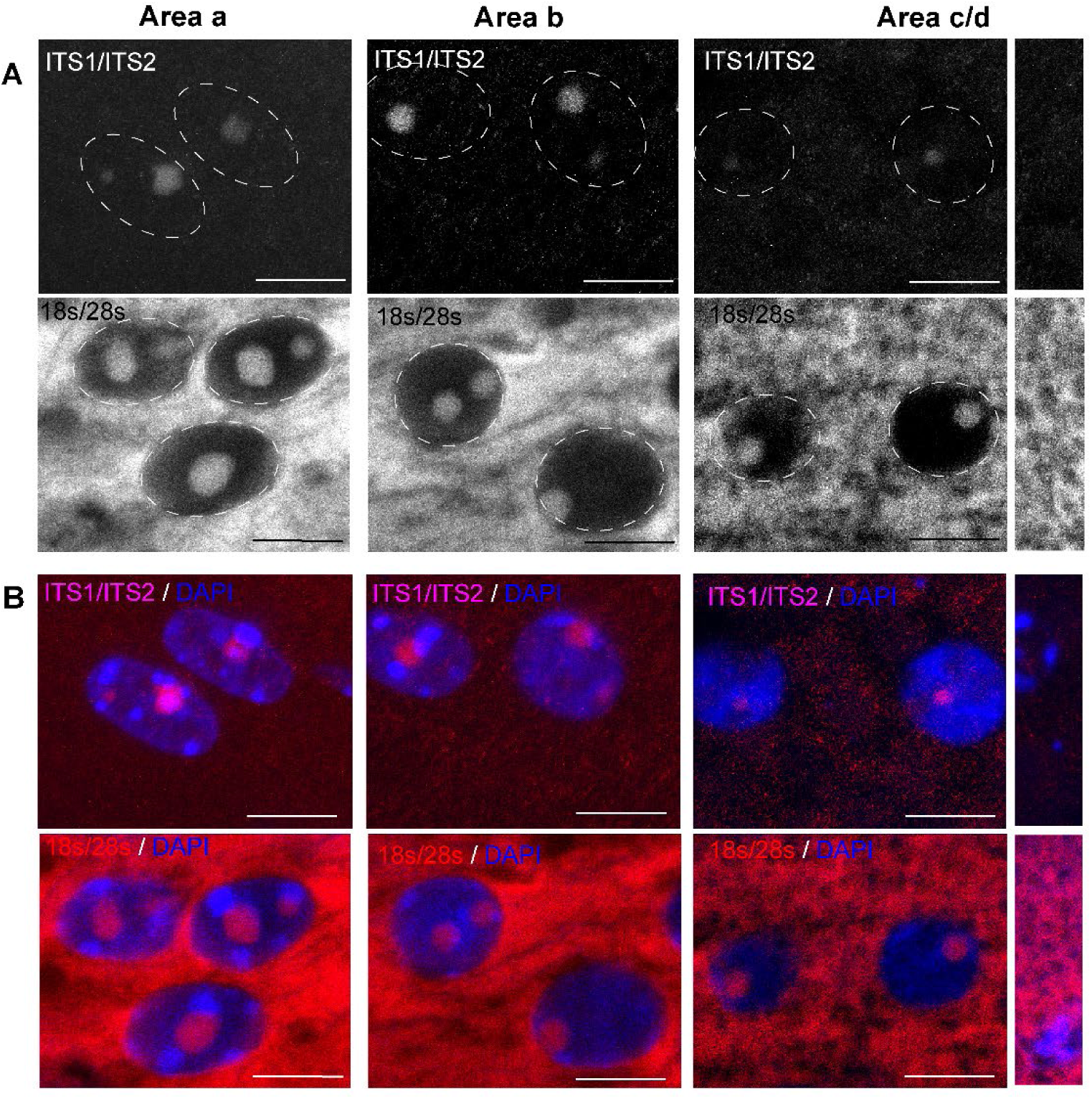
Visualization of nascent rRNA transcription throughout the lens fiber cell differentiation. (A) Black and white images using ITS1/ITS2 (top) and 18S/28S (bottom) set of probes. Nucleus delineated in dotted lines. (B) Merged images of ITS1/ITS2 probes (magenta - top) and 18S/28S probes (red - bottom) and nuclei stained with DAPI (blue) in areas a, b, c, and d. 5’ETS: 5’external transcribed spacer; 3’ETS: 3’external transcribed spacer. Scale bars = 10 µm.

**FIGURE 8.**
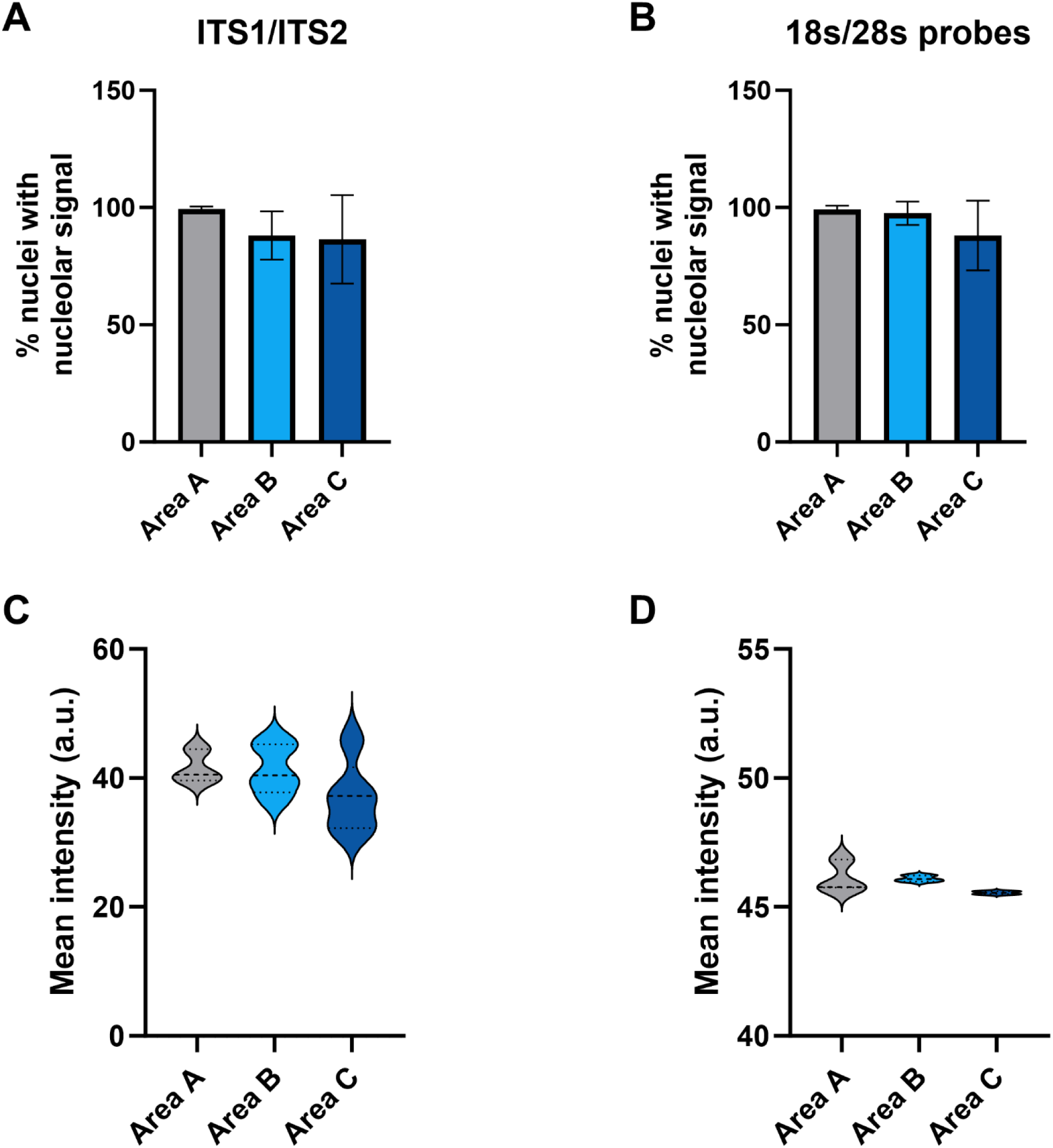
Quantification of nascent rRNAs and mature rRNAs in lens fiber cells throughout their differentiation. (A) Quantification of nuclei (in percentage) displaying nucleolar signal for ITS1/ITS2 FISH probes in areas a, b and c. (B) Quantification of nuclei (in percentage) displaying nucleolar signal for 18S/28S FISH probes. No significant differences were observed in both groups, n=3 biological replicates. (C) Under-processed rRNAs puncta’s fluorescence intensity measured as signal intensity in arbitrary units (a.u.) in areas a, b and c. (D) Mature rRNAs puncta’s fluorescence intensity measured as signal intensity in arbitrary units (a.u.) in areas a, b and c. Fluorescence intensity was calculated in deconvoluted images and are represented by arbitrary units (a.u.). No significant differences were observed, n=3 biological replicates.

Next, we quantified the nucleolar signals observed throughout areas a-c for both nascent and mature rRNAs combining signals of the ITS and 18S/28S probes. We found that areas a, b and c display comparable numbers of nascent rRNAs co-localizing with the nucleoli (Fig. 8A). Interestingly, the quantities of mature rRNAs in the nucleus of cells in areas a, b and c have also not shown significant differences (Fig. 8B). Our data also show that cells in area c display comparable numbers of both nascent and mature rRNAs within the cells in area a, at earliest stages of lens fiber cells differentiation. In addition, we quantified the mean intensity of the nuclear signals for ITS1/ITS2 and 18S/28S probes throughout areas a and c (Figs. 8C, D). Taken together, these quantitative measurements demonstrate that the levels of pre-mature and mature rRNA are maintained invariable from area a to area c in differentiating lens fiber cells.

### Transcriptional machineries are maintained prior final nuclear desintegration

The nuclear condensation and transfer of proteins into the cytoplasm (Zhao et al., 2016; Limi et al. 2018) obviously represent a major challenge for all normal nuclear functions. We thus visualized UBF (Upstream Binding Transcription Factor), transcriptionally active RNA Polymerase II as a marker of active sites of mRNA expression, and SC-35/Srsf2 protein of the nuclear speckles (Figs. 9A-C). UBF is a nucleolar DNA-binding phosphoprotein essential for RNA Polymerase I transcriptional machinery (Theophanous et al., 2023). We observed the co-localization of UBF with nucleoli of cells in areas a and b (Fig. 9B). RNA polymerase II displays a marked co-localization in nuclei of areas a and b (Fig. 9A). In area c, a few strong signals of RNA Polymerase II are detected; nevertheless, our previous studies have shown that these larger condensates represent active sites of crystallin gene expression (Limi et al. 2018, Fig. 9A). Nuclear speckles represent hubs that spatially link pre-mRNA transcription, splicing, and nuclear export (Spector and Lamond, 2011). Nuclear speckles also co-localize with nucleoli in areas a and b (Fig. 9C). Remarkably, nuclei of area c display nucleoli positive staining for all three factors: UBF, RNA Polymerase II and SC-35 (Figs. 9A-C). Taken together, these data coupled with the positive nucleolar signals of rRNA probes (Fig. 6) demonstrate that lens fiber cells are actively producing rRNAs in the nucleoli of highly condensed nuclei just prior to their denucleation.

**FIGURE 9.**
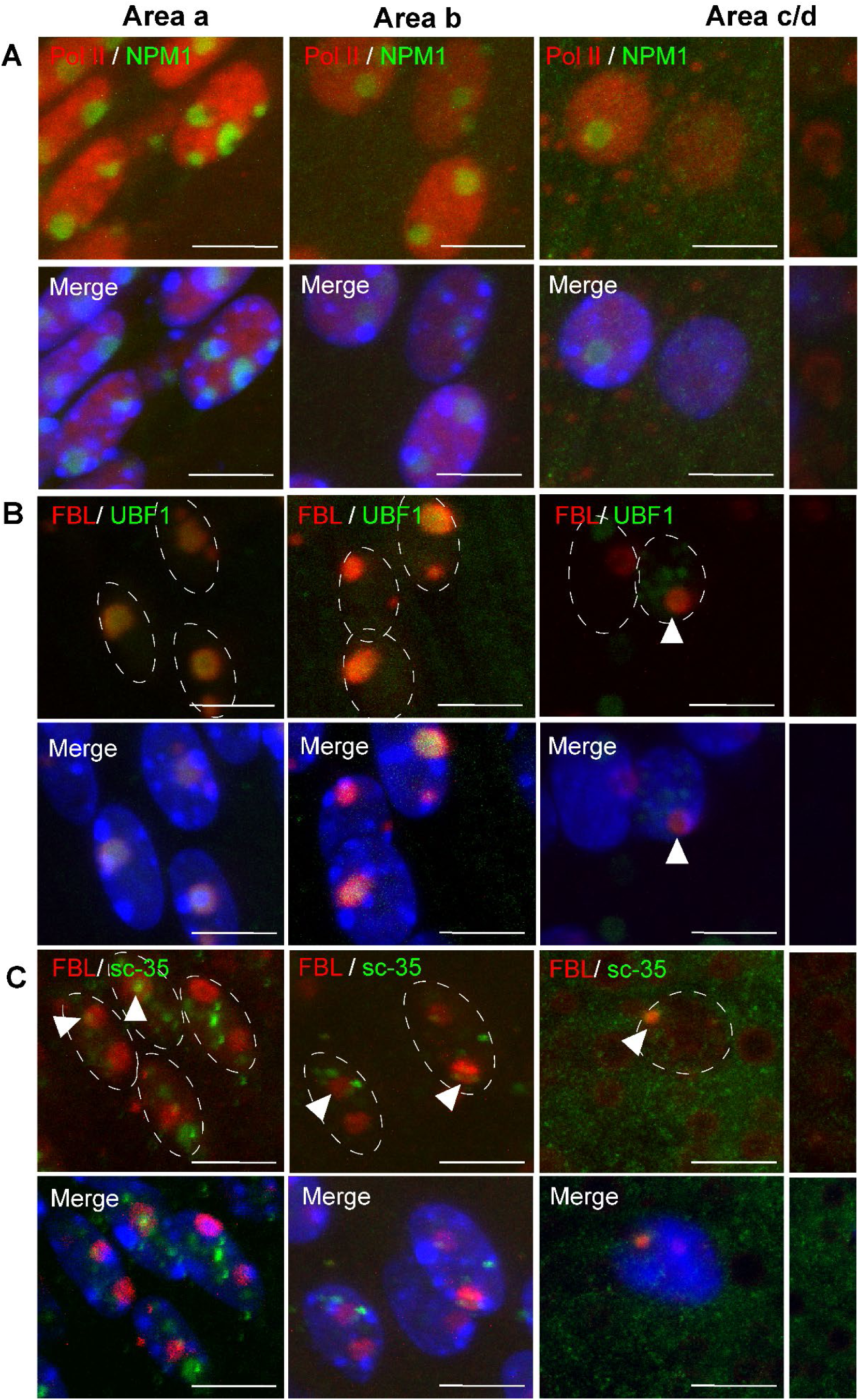
Nucleolar machinery is maintained prior final nuclear disintegration (P0.5). (A) Transcriptionally active RNA Polymerase II (Pol II, red) signals are found in the nucleoplasm outside of the nucleoli in all the areas a, b and c. Nucleolar marker nucleophosmin (NPM1) shown in green. (B) The RNA Polymerase I-specific UBF proteins (green) co-localize with nucleoli immunolabelled with fibrillarin (red) in areas a, b and c (white arrowhead). Nuclei shown in dotted white lines – top. (C) The nuclear speckles detected via Sc-35 (green) signal are observed in the nucleoli immunolabelled with fibrillarin (red) within all the areas a, b, and c (white arrowhead). Nuclei shown in dotted white lines – top. Scale bars = 10 µm. Merge images with DAPI-stained nuclei (blue).

Finally, to confirm nucleolar integrity prior to denucleation, we immunolabelled nucleolin (Anti-NCL) and nucleophosmin (Anti-NMP1), two essential nucleolar proteins that are required for ribosome biogenesis (Scott and Oeffinger, 2016). Both nucleolar proteins are observed in fiber cells nuclei in areas a and b, co-localizing with the nucleolus observed by DAPI counterstaining (Figure 10A, B). We have confirmed that fiber cells in area c also show clear intranuclear staining for nucleolin and nucleophosmin (Figs. 10A, B).

**FIGURE 10.**
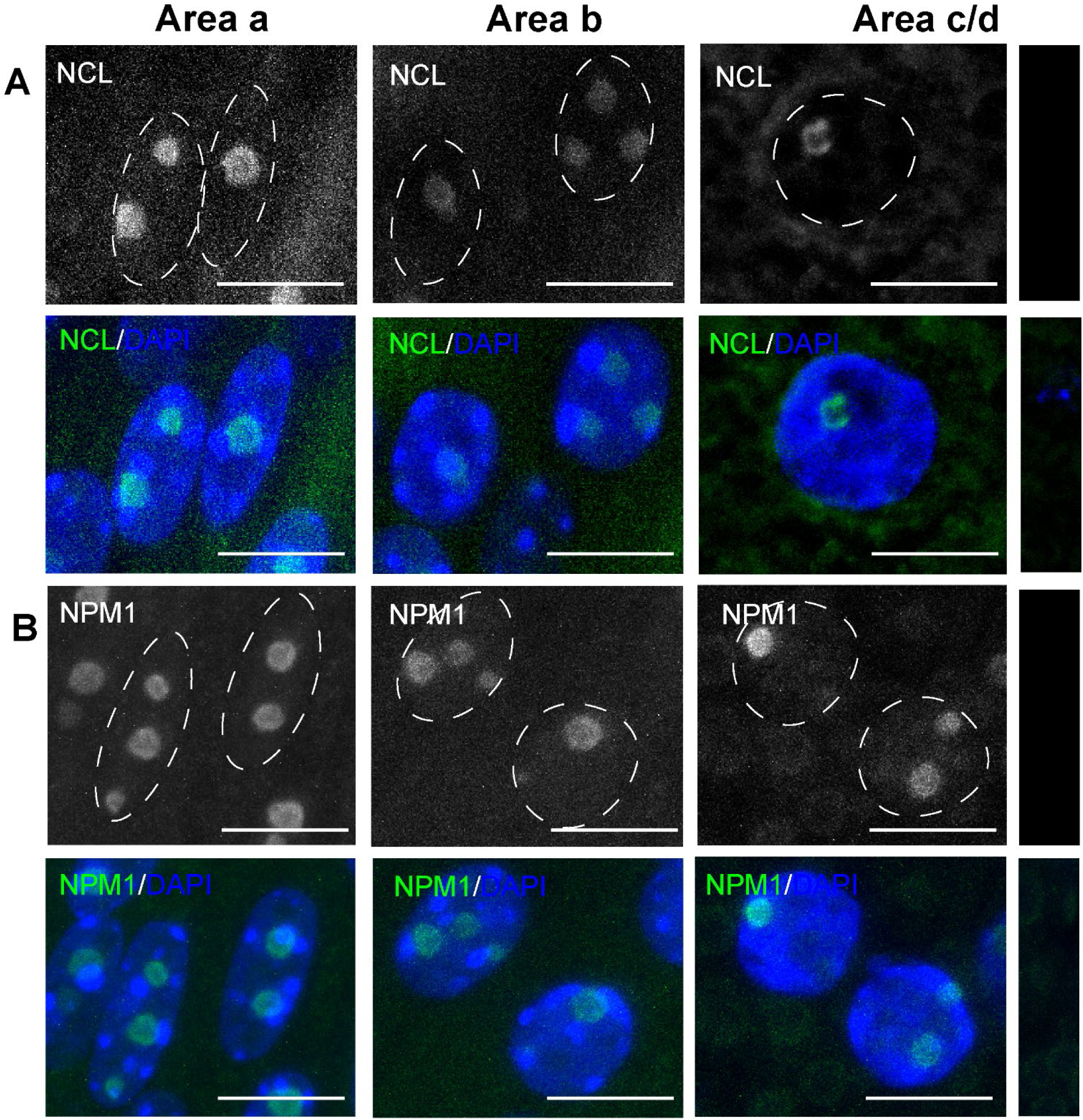
Essential nucleolar markers are present in fiber cells nuclei undergoing denucleation. (A) Nucleolin (NCL) staining in newborn lens in areas a, b and c. Black and white NCL staining, nuclei shown in dotted lines (top); NCL merged with DAPI (bottom). (B) Nucleophosmin (NPM1) staining in newborn lens in area a, b and c, nuclei shown in white dotted lines (top); NPM1 merged with DAPI (bottom). Scale bars = 10 µm. DAPI-stained nuclei (blue).

## DISCUSSION

The main goals of the present study were to evaluate how nuclear and nucleolar morphology changes throughout the highly organized temporal-spatial patterns of lens fiber cell differentiation and to determine whether rDNA transcription catalyzed by RNA Polymerase I is terminated or retained throughout the entire lifespan of lens fiber cell nuclei. We thus analyzed for the first time multiple nuclear and nucleolar markers and employed specific rRNA probes to visualize nascent and mature ribosomal RNAs throughout all stages of primary and secondary mouse lens fibers cell differentiation. The present data (Figs. 6-8) clearly show that nascent rDNA transcription persists in lens fiber cell nuclei that are located next to the gradually expanding OFZ.

Mouse genetic studies of lens fiber cell denucleation revealed initially requirements for lipoxygenase pathway enzyme Alox15 (van Leyen et al., 1998) and lens-specific acidic DNase IIβ (Nishimoto et al. 2003) co-localizing with Lamp-1 in the lysosomes in a relatively narrow region adjacent to the OFZ (Nakahara et al., 2007). Following disassembly of the nuclear membrane, these lysosomes gain access to and degrade chromatin DNA (Bassnett, 2009). Their nuclear entry requires cell cycle regulatory kinase Cdk1 to phosphorylate lamins A and C as well as nuclear mitotic apparatus protein 1 (Numa1) (Chaffee et al., 2014). Subsequent studies found additional genes involved in these and other pathways. For example, DNA-binding transcription factors Hsf4 (Fujimoto et al., 2004), Gata3 (Maeda et al., 2009; Martynova et al., 2019) and Pax6 (He et al., 2016) directly regulate expression of the *Dnase2b* gene. The denucleation process is also disrupted following inhibition of autophagy in chick lenses (Basu et al. 2014). Retention of nuclei was also found following lens-specific depletion of ATP-dependent chromatin remodeling enzymes Brg1/Smarca4 (He et al., 2010) and Snf2h/Smarca5 (He et al., 2016) even though these enzymes are transferred into the cytoplasm in advanced lens fiber cells (Limi et al., 2018). Interestingly, depletions of DNA repair and associated proteins Ddb1 (Cang et al., 2006), Nbs1/Nbn (Yang et al., 2006), and Ncoa6 (Wang et al., 2010) also inhibit lens fiber cell denucleation suggesting “repurposing” of at least three proteins involved in DNA repair to participate in the denucleation process while active DNA repair is evidenced from the γH2AX stainings (Fig. 5).

The temporally and spatially regulated gradual changes in nuclear morphology are ideally viewed at P0.5 with already established OFZ in the central portion of the lens fiber cell compartment (Figs. 1B, C). The spherical and ovoid nuclear shapes are most common nuclear morphologies and nuclear rounding and transfer of proteins into the cytoplasm are shared between lens fiber cells and erythrocytes (Zhao et al., 2016; Limi et al., 2018). It has been proposed that nuclear shape and location within the cell are both affected by cellular morphology linked to mechanical factors/constraints and driven by chromatin reorganization (Webster et al., 2009). From the first point of view, the hexagonal-shaped lens fiber cells are unique given their length (cell length elongation factor ∼1,000-fold) and mechanical properties related to the lens accommodation (Bassnett et al., 2011; Cheng et al., 2017; Cheng et al., 2019); nevertheless, their nuclei appear highly organized within the lens fiber cell cytoplasm (Fig. 1). From the point of view of chromatin organization, nuclear condensation is marked by a small number of large molecular condensates of transcriptionally active RNA Polymerase II that co-localize with nascent mRNA transcription of multiple crystallin genes expressed at their maximal levels (Limi et al., 2018). Most recently, our studies of 3D-nuclear organization of lens fiber cells compared to the lens epithelium show major changes in chromatin looping of many loci encoding lens fiber cell structural proteins (Camerino et al., 2024). In addition, there is a remarkable redistribution of DNA-binding CTCF proteins, major organizers of chromatin looping (Ghirlando and Felsenfeld, 2016). In the lens fiber cell chromatin, ChIP-seq data show that CTCF proteins are localized along the RNA Polymerase II (Camerino et al., 2024).

The present studies provide new insights into the very high transcriptional-translation outputs of lens fiber cell nuclei particularly regarding crystallin proteins that represent as much as 90% of lens water-soluble protein fraction (Bassnett et al., 2011). However, multiple physiological limits exist for these processes, including sub-optimal delivery of nutrients to the lens due to the hyaloid vascular system regression (Lang, 1997; McKeller et al. 2002; Chen et al., 2008), highly hypoxic conditions in the lens fiber cell compartment (Shui and Beebe, 2008; Brennan et al. 2020), and generation of the OFZ, including abrupt nuclear degradation. Since the rRNA production highly exceeds quantities of mRNAs it is possible that the lens fiber cells had to evolve a delicate system to find a compromise between these resource-demanding processes related to the RNA Polymerase I system (Grummt and Ladurner, 2008; Cerqueira and Lemos, 2019, Daiß et al., 2023). One possibility is to reduce expression of rRNAs to boost production of crystallin mRNAs, or to reduce crystallin gene nascent transcription in favor of rRNA to generate additional ribosomes. The present data clearly demonstrate that the lens fiber cell nuclei, approaching their degradation, are capable of high transcriptional outputs by both RNA Polymerase I and II systems. This resilience also provides indirect evidence about the functionality of molecular condensates (Brangwynne et al., 2011; Banani et al., 2017; Sharp et al., 2021) within the lens fiber cell nuclei approaching their physical end. Thus, our data suggest that even though the resources within the lens fiber cells are limited at this developmental stage, the production of rRNA and mRNA remain active till the very end of the nuclear integrity.

In conclusion, the present studies demonstrate that lens fiber cell differentiation and denucleation is a unique model to study basic transcriptional mechanisms and chromatin organization under various highly restrictive conditions. The process of terminal denucleation will require additional mechanistic insights using a set of fluorescently marked lysosomal, nucleoplasmic, and membrane proteins followed by their *in vivo* imaging. Interestingly, the translational mechanisms in both the maturing and denucleated lens fiber cells remain also to be established as evidence exist about differential abundances of ribosomal proteins between the lens epithelium and lens fibers (Zhao et al. 2019). Thus, follow up studies of ribosomal heterogeneity and plasticity in various model systems (Trahan and Oeffinger, 2023) are well justified in future studies of protein translation during lens fiber cell elongation and maturation.

## MATERIALS AND METHODS

### Mice and tissue collection

Mice husbandry and tissue collection were conducted in accordance with the approved protocol of the Albert Einstein College of Medicine Animal Institute Committee and the ARVO statement for the use of animals in eye research. The day of confirmed vaginal plug was considered as embryonic stage E0.5. Individual eyes were harvested from embryonic day E14.5, E16.5, E18.5 and newborn (P0) CD1 mice. Females were euthanized by CO_2_, mouse embryos were collected and eyes dissected. Postnatal mice were euthanized by CO_2_ followed by decapitation and eyeballs were dissected. Tissues were fixed overnight in 4% paraformaldehyde at 4° C followed by cryoprotection in 30% sucrose overnight at 4° C and embedded in optimal cutting temperature (OCT).

### Immunofluorescence

Slides were kept at room temperature for 1hour followed by antigenic retrieval for 10 minutes at 90 degrees in citric acid solution (H3300, Vector Labs) following manufacturer instructions. Slides were cooled down for 10 minutes, and blocked in 10% NGS (50062Z, Thermo Fisher), 1% BSA (A4737-25G, Sigma), 0.1% Triton (9036-19-5, Sigma) in PBS for 1 hour at room temperature. Slides were subsequently incubated (1:100) in blocking solution at 4^0^C with primary antibodies: anti-Fibrillarin (NB300-269, Novus Biologicals); anti-Lamin B (66095-1-Ig, Proteintech); anti-γH2AX (sc-377452, Santa Cruz); anti-RNA Polymerase II (ab26721, Abcam); anti-UBF1 (ab244287, Abcam); anti-SC35 (sc-53518, Santa Cruz); anti-NCL (10556-1-AP, Proteintech), and anti-NPM1 (32-5200, Thermo). After overnight incubation, slides were washed three times in PBS and incubated for 1h at room temperature with secondary antibodies Goat anti-Rabbit IgG Alexa Fluor™ Plus 555 (A32732, Thermo) or Goat anti-Mouse IgG, Alexa Fluor™ 555 (A-21422, Thermo). Sections were washed three times in PBS, incubated 5 minutes at room temperature with DAPI and mounted in ProLong™ Diamond Antifade Mountant (P36961, Thermo).

### Fluorescence *in situ* hybridization (FISH)

The procedure to generate probes for ribosomal RNA and imaging is described elsewhere (Qian et al., 2006; Chebrout et al., 2022). Probes located in different regions of the 47S pre-RNA were used as described above (Chebrout et al., 2022). Internal-transcribed spacers 1 and 2 (ITS1 and ITS2) enable the localization of under-processed rRNAs, whereas probes for the 18S and 28S regions localize mature pre-RNAs in the nucleus and/or cytoplasm. rRNA probes oligos were designed from the mouse pre-rRNA gene (ITS1: 3’-TAG-ACA-CGG-AAG-AGC-CGG-ACG-GGA-AAG-A-5’-Cy3; 3’-ITS2: CCA-GCG-CAA-GAC-CCA-AAC-ACA-CAC-AGA-5’-Cy3; 18S: 3’-CCA-TTA-TTC-CTA-GCT-GCG-GTA-TCC-AGG-CGG-5’-Cy5; 28S: 3’-GAG-GGA-ACC-AGC-TAC-TAG-ATG-GTT-CGA-TTA-5’-Cy5). Slides were immersed in Reveal Deckloaking Buffer (RV1000, Biocare Medical) at 90^0^C for 5 minutes. Subsequently, slides were processed through several treatments to reduce tissue autofluorescence (Limi et al., 2019), and incubated in 50% formamide pre-hybridization buffer (50% formamide in 20x SSC Buffer) at 37° for 1 hour. Cryosections were hybridized with 125nM ITS1/ITS2 or 18S/28S probes overnight in hybridization buffer (10% dextrane sulfate, 20x SSC, 50% Formamide, 10mg/ml E. coli tRNA, 200mM VRC, 20mg/ml BSA) at 37^0^ C. Post-hybridization washes with pre-hybridization buffer were conducted, slides were incubated with DAPI for 15 minutes and mounted in ProLong™ Diamond Antifade Mountant (P36961, Thermo).

### Lens segmentation, imaging, quantification, statistical methods, and analyses

To analyze the progression of lens fiber cells differentiation, lens tissue was symmetrically divided into four regions from the periphery to the center labelled as areas a, b, c, and d. Area a comprises early differentiated fiber cells, area b intermediate fiber cells and, area c advanced fiber cells prior to the denucleation. Area d represents disintegrating nuclei, and the organelle free zone is only observed in newborn lenses (Limi et al., 2019). Nuclear rRNA were observed using Zeiss AxioObserver CLEM microscope. Three-dimensional image data were acquired using the Leica SP8 Confocal at x63 oil-immersion objective plus 2.0 zoom factor. Cell quantification and number of nuclei with nucleolar rRNA signal was conducted using ImageJ software (http://imagej.nih.gov/ij/; NIH) throughout the entire z-stack section of each image. Images were deconvoluted in Volocity software (PerkinElmer) prior to fluorescence intensity analysis, indicated by “mean intensity (a.u.)” as we previously described (Limi et al., 2019). At least 50 nuclei from n=3 animals were analyzed per segmented area. Note smaller number of nuclei in zone d is due to their degradation. Unpaired t test or One-way ANOVA with multiple comparisons posttest was performed using GraphPad Prism 10 for Windows (www.graphpad.com; GraphPad software) and statistical significance was considered according to p<0.001***; p<0.01**; p<0.5*.

## ACKNOWLEDGEMENTS

We thank Dr. Noura Ghazale for her helpful insights on the *in-situ* hybridization protocol. We thank the Analytical Imaging Facility (AIF) at the Albert Einstein College of Medicine for their help with microscopy and analysis.

## ABBREVIATIONS

DFC: Dense fibrillar component
ER: endoplasmic reticulum
FC: fibrillar center
FISH: fluorescence *in situ* hybridization
GC: granular component
OFZ: organelle-free zone
3D: 3-dimensional

---

**Supplementary Figure S1. Nucleolar counting by area in newborn and embryonic lens.** (A) Newborn lens fiber cells display an equal number of nucleoli per nuclei in areas A, B and C. Most of the cells analyzed display one single nucleolus per nucleus. (B) Embryonic lens fiber cells display similar numbers of nucleoli per nuclei in areas a, b, c and d.

## REFERENCES

Banani SF, Lee HO, Hyman AA, Rosen MK. 2017. Biomolecular condensates: organizers of cellular biochemistry. 2017. Nat Rev Mol Cell Biol 18: 285–298. doi: 10.1038/nrm.2017.7

Bassnett S. 2009. On the mechanism of organelle degradation in the vertebrate lens. Exp Eye Res 88: 133–9. doi: 10.1016/j.exer.2008.08.017

Bassnett S, Shi Y, Vrensen GF. 2011. Biological glass: structural determinants of eye lens transparency. Philos Trans R Soc Lond B Biol Sci 366: 1250–64. doi: 10.1098/rstb.2010.0302

Basu S, Rajakaruna S, Reyes B, Van Bockstaele E, Menko AS. 2014. Suppression of MAPK/JNK-MTORC1 signaling leads to premature loss of organelles and nuclei by autophagy during terminal differentiation of lens fiber cells. Autophagy 10: 1193–211. doi: 10.4161/auto.28768

Beebe DC. 2008. Maintaining transparency: a review of the developmental physiology and pathophysiology of two avascular tissues. Semin Cell Dev Biol 19: 125–33. doi: 10.1016/j.semcdb.2007.08.014

Boisvert FM, van Koningsbruggen S, Navascués J, Lamond AI. 2007. The multifunctional nucleolus. Nat Rev Mol Cell Biol 8: 574–85. doi: 10.1038/nrm2184

Brangwynne CP, Mitchison TJ, Hyman AA. 2011. Active liquid-like behavior of nucleoli determines their size and shape in *Xenopus laevis* oocytes. Proc Natl Acad Sci U S A 108: 4334–9. doi: 10.1073/pnas.1017150108

Brennan LA, McGreal-Estrada R, Logan CM, Cvekl A, Menko AS, Kantorow M. 2018. BNIP3L/NIX is required for elimination of mitochondria, endoplasmic reticulum and Golgi apparatus during eye lens organelle-free zone formation. Exp Eye Res. 174: 173–184. doi: 10.1016/j.exer.2018.06.003

Brennan L, Disatham J, Kantorow M. 2020. Hypoxia regulates the degradation of non-nuclear organelles during lens differentiation through activation of HIF1a. Exp Eye Res. 198:108129. doi: 10.1016/j.exer.2020.108129.

Brennan L, Disatham J, Kantorow M. 2021. Mechanisms of organelle elimination for lens development and differentiation. Exp Eye Res 209: 108682. doi: 10.1016/j.exer.2021.108682

Camerino M, Chang W, Cvekl A. 2024. Analysis of long-range chromatin contacts, compartments and looping between mouse embryonic stem cells, lens epithelium and lens fibers. Epigenetics Chromatin 17: 10. doi: 10.1186/s13072-024-00533-x

Cang Y, Zhang J, Nicholas SA, Bastien J, Li B, Zhou P, Goff SP. 2006. Deletion of DDB1 in mouse brain and lens leads to p53-dependent elimination of proliferating cells. Cell 127: 929–40. doi: 10.1016/j.cell.2006.09.045

Cech TR, Steitz JA. 2014. The noncoding RNA revolution-trashing old rules to forge new ones. Cell 157:77–94. doi: 10.1016/j.cell.2014.03.008.

Cerqueira AV, Lemos B. 2019. Ribosomal DNA and the Nucleolus as Keystones of Nuclear Architecture, Organization, and Function. Trends Genet 35: 710–723. doi: 10.1016/j.tig.2019.07.011

Chaffee BR, Shang F, Chang ML, Clement TM, Eddy EM, Wagner BD, Nakahara M, Nagata S, Robinson ML, Taylor A. 2014. Nuclear removal during terminal lens fiber cell differentiation requires CDK1 activity: appropriating mitosis-related nuclear disassembly. Development 141: 3388–98. doi: 10.1242/dev.106005

Chebrout M, Koné MC, Jan HU, Cournut M, Letheule M, Fleurot R, Aguirre-Lavin T, Peynot N, Jouneau A, Beaujean N, Bonnet-Garnier A. 2022.Transcription of rRNA in early mouse embryos promotes chromatin reorganization and expression of major satellite repeats. J Cell Sci 135: jcs258798. doi: 10.1242/jcs.258798

Chen Y, Doughman YQ, Gu S, Jarrell A, Aota S, Cvekl A, Watanabe M, Dunwoodie SL, Johnson RS, et al. 2008. Cited2 is required for the proper formation of the hyaloid vasculature and for lens morphogenesis. Development 135: 2939–48 doi: 10.1242/dev.021097

Cheng C, Nowak RB, Fowler VM. 2017. The lens actin filament cytoskeleton: Diverse structures for complex functions. Exp Eye Res 156: 58–71. doi: 10.1016/j.exer.2016.03.005

Cheng C, Parreno J, Nowak RB, Biswas SK, Wang K, Hoshino M, Uesugi K, Yagi N, Moncaster JA, Lo WK, Pierscionek B, Fowler VM. 2019. Age-related changes in eye lens biomechanics, morphology, refractive index and transparency. Aging (Albany NY) 11: 12497–12531. doi: 10.18632/aging.102584

Correll CC, Bartek J, Dundr M. 2019.The Nucleolus: A Multiphase Condensate Balancing Ribosome Synthesis and Translational Capacity in Health, Aging and Ribosomopathies. Cells 8: 869. doi: 10.3390/cells8080869

Cvekl A, Zhang X. 2017. Signaling and Gene Regulatory Networks in Mammalian Lens Development. Trends Genet 33: 677–702. doi: 10.1016/j.tig.2017.08.001

Cvekl A, Eliscovich C. 2021. Crystallin gene expression: Insights from studies of transcriptional bursting. Exp Eye Res 207: 108564. doi: 10.1016/j.exer.2021.108564

Dahm R, Gribbon C, Quinlan RA, Prescott AR. 1998. Changes in the nucleolar and coiled body compartments precede lamina and chromatin reorganization during fibre cell denucleation in the bovine lens. Eur J Cell Biol 75: 237–46. doi: 10.1016/S0171-9335(98)80118-0

Daiß JL, Griesenbeck J, Tschochner H, Engel C. 2023. Synthesis of the ribosomal RNA precursor in human cells: mechanisms, factors and regulation. Biol Chem 404: 1003–1023. doi: 10.1515/hsz-2023-0214

Dundr M, Misteli T. 2001. Functional architecture in the cell nucleus. Biochem J. 356:297–310. doi: 10.1042/0264-6021:3560297.

Feng S, Manley JL. 2022. Beyond rRNA: nucleolar transcription generates a complex network of RNAs with multiple roles in maintaining cellular homeostasis. Genes Dev. 36:876–886. doi: 10.1101/gad.349969.122.

Fujimoto M, Izu H, Seki K, Fukuda K, Nishida T, Yamada S, Kato K, Yonemura S, Inouye S, Nakai A. 2004. HSF4 is required for normal cell growth and differentiation during mouse lens development. EMBO J 23: 4297–306. doi: 10.1038/sj.emboj.7600435

Granneman, S, Baserga, SJ. 2005. Crosstalk in gene expression: coupling and co-regulation of rDNA transcription, pre-ribosome assembly and pre-rRNA processing. Curr Opin Cell Biol 17: 281–6. doi: 10.1016/j.ceb.2005.04.001

Gheyas R, Menko AS. 2023. The involvement of caspases in the process of nuclear removal during lens fiber cell differentiation. Cell Death Discov 9: 386. doi: 10.1038/s41420-023-01680-y

Ghirlando R, Felsenfeld G. 2016. CTCF: making the right connections. Genes Dev 30: 881–91. doi: 10.1101/gad.277863.116

Gribbon C, Dahm R, Prescott AR, Quinlan RA. 2002. Association of the nuclear matrix component NuMA with the Cajal body and nuclear speckle compartments during transitions in transcriptional activity in lens cell differentiation. Eur J Cell Biol. 81: 557–66. doi: 10.1078/0171-9335-00275

Grummt I, Ladurner AG. 2008. A metabolic throttle regulates the epigenetic state of rDNA. Cell 133: 577–80. doi: 10.1016/j.cell.2008.04.026

Grummt I. 2003. Life on a planet of its own: regulation of RNA polymerase I transcription in the nucleolus. Genes Dev. 17: 1691–702. doi: 10.1101/gad.1098503R

He S, Pirity MK, Wang WL, Wolf L, Chauhan BK, Cveklova K, Tamm ER, Ashery-Padan R, Metzger D, Nakai A, Chambon P, Zavadil J, Cvekl A. 2010. Chromatin remodeling enzyme Brg1 is required for mouse lens fiber cell terminal differentiation and its denucleation. Epigenetics Chromatin 3: 21. doi: 10.1186/1756-8935-3-21

He S, Limi S, McGreal RS, Xie Q, Brennan LA, Kantorow WL, Kokavec J, Majumdar R, Hou H Jr, Edelmann W, Liu W, Ashery-Padan R, Zavadil J, Kantorow M, Skoultchi AI, Stopka T, Cvekl A. 2016. Chromatin remodeling enzyme Snf2h regulates embryonic lens differentiation and denucleation. Development 143: 1937–47. doi: 10.1242/dev.135285

Hori Y, Engel C, Kobayashi T. 2023. Regulation of ribosomal RNA gene copy number, transcription and nucleolus organization in eukaryotes. Nat Rev Mol Cell Biol 24: 414–429. doi: 10.1038/s41580-022-00573-9

Lafontaine DLJ, Riback JA, Bascetin R, Brangwynne CP. 2021.The nucleolus as a multiphase liquid condensate. Nat Rev Mol Cell Biol 22: 165–182. doi: 10.1038/s41580-020-0272-6

Lang RA. 1997. Apoptosis in mammalian eye development: lens morphogenesis, vascular regression and immune privilege. Cell Death Differ 4: 12–20. doi: 10.1038/sj.cdd.4400211

Limi S, Senecal A, Coleman R, Lopez-Jones M, Guo P, Polumbo C, Singer RH, Skoultchi AI, Cvekl A. 2018. Transcriptional burst fraction and size dynamics during lens fiber cell differentiation and detailed insights into the denucleation process. J Biol Chem 293: 13176–13190. doi:10.1074/jbc.RA118.001927

Limi S, Zhao Y, Guo P, Lopez-Jones M, Zheng D, Singer RH, Skoultchi AI, Cvekl A. 2019. Bidirectional Analysis of Cryba4-Crybb1 Nascent Transcription and Nuclear Accumulation of Crybb3 mRNAs in Lens Fibers. Invest Ophthalmol Vis Sci 60: 234–244. doi: 10.1167/iovs.18-25921

Ma N, Matsunaga S, Takata H, Ono-Maniwa R, Uchiyama S, Fukui K. 2007. Nucleolin functions in nucleolus formation and chromosome congression. J Cell Sci 120: 2091–105. doi: 10.1242/jcs.008771

Maeda A, Moriguchi T, Hamada M, Kusakabe M, Fujioka Y, Nakano T, Yoh K, Lim KC, Engel JD, Takahashi S. 2009. Transcription factor GATA-3 is essential for lens development. Dev Dyn 238: 2280–91. doi: 10.1002/dvdy.22035

Martynova E, Zhao Y, Xie Q, Zheng D, Cvekl A. 2019. Transcriptomic analysis and novel insights into lens fibre cell differentiation regulated by Gata3. Open Biol. 9: 190220. doi: 10.1098/rsob.190220

McKeller RN, Fowler JL, Cunningham JJ, Warner N, Smeyne RJ, Zindy F, Skapek SX. 2002. The Arf tumor suppressor gene promotes hyaloid vascular regression during mouse eye development. Proc Natl Acad Sci U S A 99: 3848–53. doi: 10.1073/pnas.05248419

McStay B. 2016. Nucleolar organizer regions: genomic ‘dark matter’ requiring illumination. Genes Dev 30: 1598–610. doi: 10.1101/gad.283838.116

Misteli T. 2020. The Self-Organizing Genome: Principles of Genome Architecture and Function. Cell. 183:28–45. doi: 10.1016/j.cell.2020.09.014.

Nakahara M, Nagasaka A, Koike M, Uchida K, Kawane K, Uchiyama Y, Nagata S. 2007. Degradation of nuclear DNA by DNase II-like acid DNase in cortical fiber cells of mouse eye lens. FEBS J 274: 3055–64. doi: 10.1111/j.1742-4658.2007.05836.x

Ni C, Buszczak M. 2023. The homeostatic regulation of ribosome biogenesis. Semin Cell Dev Biol. 136:13–26. doi: 10.1016/j.semcdb.2022.03.043.

Nishimoto S, Kawane K, Watanabe-Fukunaga R, Fukuyama H, Ohsawa Y, Uchiyama Y, Hashida N, Ohguro N, Tano Y, Morimoto T, Fukuda Y, Nagata S. 2003. Nuclear cataract caused by a lack of DNA degradation in the mouse eye lens. Nature 424: 1071–4. doi: 10.1038/nature01895

Olson MO, Dundr M, Szebeni A. 2000. The nucleolus: an old factory with unexpected capabilities. Trends Cell Biol 10: 189–96. doi: 10.1016/s0962-8924(00)01738-4

Qian J, Lavker RM, Tseng H. 2006. Mapping ribosomal RNA transcription activity in the mouse eye. Dev Dyn 235: 1984–93. doi: 10.1002/dvdy.20827

Roeder RG. 2019. 50+ years of eukaryotic transcription: an expanding universe of factors and mechanisms. Nat Struct Mol Biol 26: 783–791. doi: 10.1038/s41594-019-0287-x

Rogerson C, Bergamaschi D, O’Shaughnessy RFL. 2018. Uncovering mechanisms of nuclear degradation in keratinocytes: A paradigm for nuclear degradation in other tissues. Nucleus 9: 56–64. doi: 10.1080/19491034.2017.1412027

Rowan S, Chang ML, Reznikov N, Taylor A. 2017. Disassembly of the lens fiber cell nucleus to create a clear lens: The p27 descent. Exp Eye Res 156: 72–78. doi: 10.1016/j.exer.2016.02.011

Scott DD, Oeffinger M. 2016. Nucleolin and nucleophosmin: nucleolar proteins with multiple functions in DNA repair. Biochem Cell Biol. 94:419–432. doi: 10.1139/bcb-2016-0068.

Sharp PA, Chakraborty AK, Henninger JE, Young RA. 2021. RNA in formation and regulation of transcriptional condensates. RNA 28: 52–57. doi: 10.1261/rna.078997.121

Shore D, Albert B. 2022. Ribosome biogenesis and the cellular energy economy. Curr Biol. 32 : R611–R617. doi: 10.1016/j.cub.2022.04.083.

Shui YB, Beebe DC. 2008. Age-Dependent Control of Lens Growth by Hypoxia. Invest Ophthalmol Vis Sci 49: 10023–1029. doi: 10.1167/iovs.07-1164

Spector DL, Lamond AI. 2011. Nuclear speckles. Cold Spring Harb Perspect Biol 3: a000646. doi: 10.1101/cshperspect.a000646

Sun J, Rockowitz S, Chauss D, Wang P, Kantorow M, Zheng D, Cvekl A. 2015. Chromatin features, RNA polymerase II and the comparative expression of lens genes encoding crystallins, transcription factors, and autophagy mediators. Mol Vis 21: 955–73. PMID: 26330747

Theophanous A, Christodoulou A, Mattheou C, Sibai DS, Moss T, Santama N. 2023. Transcription factor UBF depletion in mouse cells results in downregulation of both downstream and upstream elements of the rRNA transcription network. J Biol Chem 299: 105203. doi: 10.1016/j.jbc.2023.105203.

Trahan C, Oeffinger M. 2023. The Importance of Being RNA-est: considering RNA-mediated ribosome plasticity. RNA Biol. 20:177–185. doi: 10.1080/15476286.2023.2204581.

Van Leyen K, Duvoisin RM, Engelhardt H, Wiedmann M. 1998. A function for lipoxygenase in programmed organelle degradation. Nature 395: 392–5. doi: 10.1038/26500

Vermunt MW, Zhang D, Blobel GA. 2018. The interdependence of gene-regulatory elements and the 3D genome. J Cell Biol. 218:12–26. doi: 10.1083/jcb.201809040.

Wang WL, Li Q, Xu J, Cvekl A. 2010. Lens fiber cell differentiation and denucleation are disrupted through expression of the N-terminal nuclear receptor box of NCOA6 and result in p53-dependent and p53-independent apoptosis. Mol Biol Cell 21: 2453–68. doi: 10.1091/mbc.e09-12-1031

Webster M, Witkin KL, Cohen-Fix O. 2009. Sizing up the nucleus: nuclear shape, size and nuclear-envelope assembly. J Cell Sci 122:1477–86. doi: 10.1242/jcs.037333

White RJ. 2011. Transcription by RNA polymerase III: more complex than we thought. Nat Rev Genet 12: 459–63. doi: 10.1038/nrg3001

Wu J, Xiao J, Zhang Z, Wang X, Hu S, Yu J. 2014. Ribogenomics: the science and knowledge of RNA. Genomics Proteomics Bioinformatics 12: 57–63. doi: 10.1016/j.gpb.2014.04.002

Yang YG, Frappart PO, Frappart L, Wang ZQ, Tong WM. 2006. A novel function of DNA repair molecule Nbs1 in terminal differentiation of the lens fibre cells and cataractogenesis. DNA Repair (Amst) 5: 885–93. doi: 10.1016/j.dnarep.2006.05.004

Yoneda M, Nakagawa T, Hattori N, Ito T. 2021. The nucleolus from a liquid droplet perspective. J Biochem 170: 153–162. doi: 10.1093/jb/mvab090

Zentner GE, Balow SA, Scacheri PC. 2014. Genomic characterization of the mouse ribosomal DNA locus. G3 (Bethesda) 4: 243–54. doi: 10.1534/g3.113.009290

Zhao B, Mei Y, Schipma MJ, Roth EW, Bleher R, Rappoport JZ, Wickrema A, Yang J, Ji P. 2016. Nuclear Condensation during Mouse Erythropoiesis Requires Caspase-3-Mediated Nuclear Opening. Dev Cell 36: 498–510. doi: 10.1016/j.devcel.2016.02.001

Zhao Y, Wilmarth PA, Cheng C, Limi S, Fowler VM, Zheng D, David LL, Cvekl A. 2019. Proteome-transcriptome analysis and proteome remodeling in mouse lens epithelium and fibers. Exp Eye Res 179: 32–46. doi: 10.1016/j.exer.2018.10.011

