## Supplementary figures and images for "Continuous nucleolar ribosomal RNA synthesis in differentiating lens fiber cells until abrupt nuclear degradation required for ocular lens transparency"

### Supplemental Fig 1

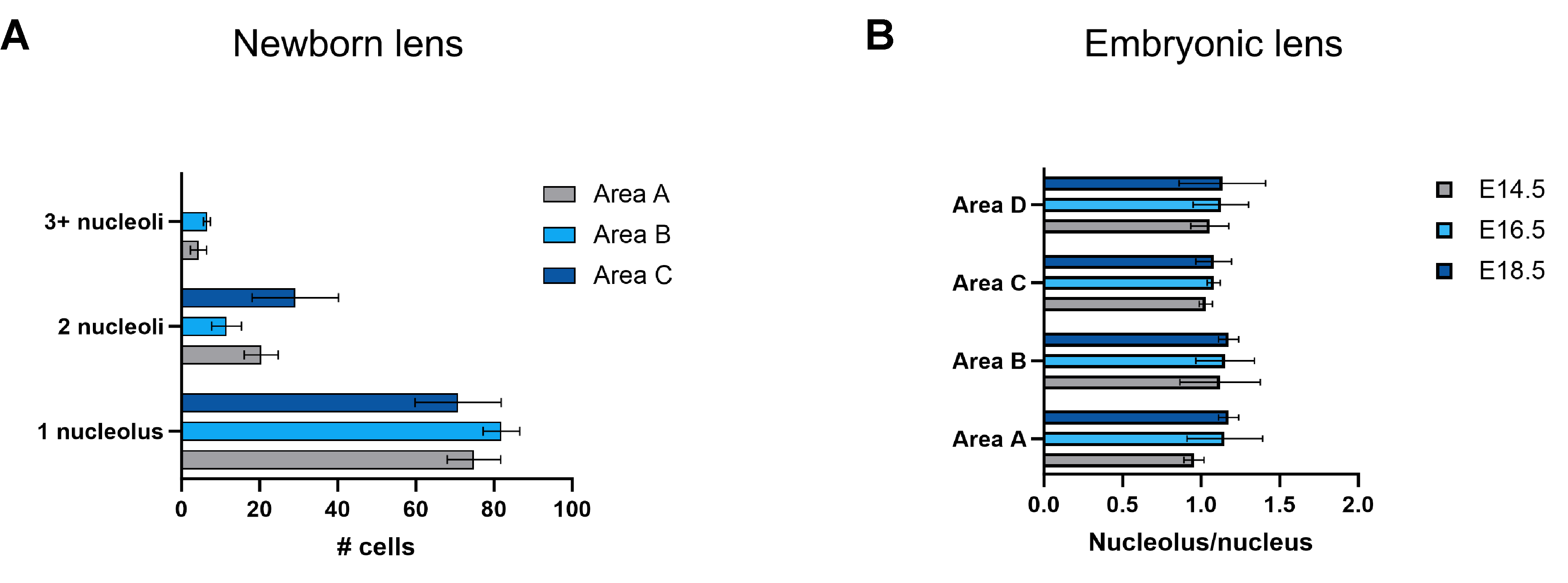
